# Chemically Accurate Relative Folding Stability of RNA Hairpins from Molecular Simulations

**DOI:** 10.1101/354332

**Authors:** Louis G. Smith, Zhen Tan, Aleksandar Spasic, Debapratim Dutta, Leslie A. Salas-Estrada, Alan Grossfield, David H. Mathews

## Abstract

This study describes a comparison between melts and simulated stabilities of the same RNAs that could be used to benchmark RNA force fields, and potentially to determine future melting experiments. Using umbrella sampling molecular simulations of three 12-nucleotide RNA hairpin stem loops, for which there are experimentally determined free energies of unfolding, we projected unfolding onto the reaction coordinate of end to end (5′ to 3′ hydroxyl oxygen) distance. We estimate the free energy change of the transition from the native conformation to a fully extended conformation—the stretched state—with no hydrogen bonds between non-neighboring bases. Each simulation was performed four times using the AMBER FF99+bsc0+χ_OL3_ force field and each window, spaced at 1 Å intervals, was sampled for 1 μs, for a total of 552 μs of simulation. We compared differences in the simulated free energy changes to analogous differences in free energies from optical melting experiments using thermodynamic cycles where the free energy change between stretched and random coil sequences is assumed to be sequence independent. The differences between experimental and simulated ΔΔ*G*° are on average 1.00 ± 0.66 kcal/mol, which is chemically accurate and suggests analogous simulations could be used predictively. We also report a novel method to identify where replica free energies diverge along the reaction coordinate, thus indicating where additional sampling would most improve convergence. We conclude by discussing methods to more economically perform such simulations.

## Introduction

RNA is central to life.^1^ It has fundamental roles in the expression of genetic information as proteins, and also functions directly in many other processes as non-coding RNA.^2–5^ It is also a source of variation and mechanism of control in cellular processes.^6–14^ Molecular interactions are the first principles of biology. Hence, all-atom simulations of RNA provide a principled framework for developing hypotheses about and interpreting experiments involving this essential biopolymer.^15–19^

All-atom simulations provide detailed information about the conformations sampled by molecules, but it can be difficult to compare simulations to experiments.^20^ Molecular dynamics (MD) uses classical mechanics to model the motions of molecules. Snapshots of the simulated system recorded periodically during the simulation approximate the molecule’s thermodynamic conformational ensemble that, if this sampling is thorough, can be compared to experimental measurements. Making this comparison, however, requires knowing the relationship between the coordinates in the snapshots and the experimental quantity measured.^21–25^

Molecular dynamics simulations require an energy model known as a force field, which estimates the potential energy from the molecular conformation. The force field functional form and parameters are approximations of the quantum mechanics underlying the molecular interactions.^26–28^ There are a number of known limitations to the RNA force fields that lead to significant artifacts in the simulations when sampling is thorough. For the commonly used AMBER force field, FF99+bsc0+χ_OL3_,^29–31^ it has been observed that in simulations stable tetraloops do not retain a folded structure,^32,33^ tetramers often miss contacts observed in proton NMR spectra and exhibit close contacts not observed in the spectra,^24,32^ internal loops do not produce ratios of major and minor conformers similar to solution structures,^34,35^ and some RNA-protein complexes are not stable for more than a microsecond.^36^ It is widely believed that no available force field cheap enough to permit adequate sampling is accurate enough to fold structured RNA *de novo*.^26,27,37^ The CHARMM 36 force field,^38^ and a revised Amber force field^39^ also do not correctly stabilize the native tetraloop structure nor accurately model tetramers.^32^ Our recent work re-parameterizing back-bone dihedrals improves simulation accuracy for tetramers and internal loops, but not tetraloops.^40^ Another revision of the AMBER force field and a new AMOEBA force field for nucleic acids were recently published, but remain to be tested by the field more broadly.^41,42^ Here we provide a different framework for testing force fields and apply it to AMBER’s FF99+bsc0+χ_OL3_ force field.

One strategy for force field benchmarking is to compare reference data to thermodynamic observables that can be simulated. One such observable with extensive reference data is the unfolding free energy change of small structured RNAs.^34,43^ Benchmarking against free energy changes tests whether the simulated volume of the RNA’s native conformational basin is accurate, in addition to testing whether it correctly represents the favorability of native contacts relative to those present in the unfolded state. This contrasts to comparisons between simulations and native structures determined by NMR or crystallography, where it is difficult to assess whether the observed conformational flexibility is consistent with the structure unless the native conformation is catastrophically lost. Simulated free energy changes have been previously compared to *in vitro* reference data for helical stacking free energy,^44–48^ stacking of modified nucleotides,^49^ small hairpin stability,^38,50–53^ and stacking of GC vs isoG-isoC pairs.^54^

This study compares the free energy change of stretching an RNA *in silico* to its free energy change of melting determined by optical melting experiments using a thermodynamic cycle.^51^ Simulating the optical melting transition with good statistics is challenging because, although strategies exist for simulating a melting event, the random coil ensemble of an RNA has a huge volume in phase space.^55,56^ Thus to focus our sampling, we simulate a transition along a distance from folded to stretched, in imitation of force denaturation by optical tweezers, hereafter referred to as pulling experiments.^51,57,58^

To simulate this stretching transition we performed umbrella sampling along the 5′ to 3′ hydroxyl oxygen distance (the end to end distance). We performed these simulations in quadruplicate for three 12-nucleotide RNAs with known free energies of unfolding and with representative NMR structures (Fig. 1). We compared the simulated change in free energy of stretching to experimentally-determined free energy changes of melting for pairs of sequences with thermody-namic cycles.^51^ The mean absolute difference between the reference and simulated ΔΔ*G*° is 0.98 kcal/mol and the propagated uncertainty in this difference is 1.16 kcal/mol. Our analysis suggests that the residuals for each sequence, which on average have an error of 0.66 kcal/mol (approximately *k_B_T* at 37°C), are accurate and precise enough to have predictive value, and set a new standard for precision in such simulations.

## Methods

### Starting Structures

**Figure 1:**
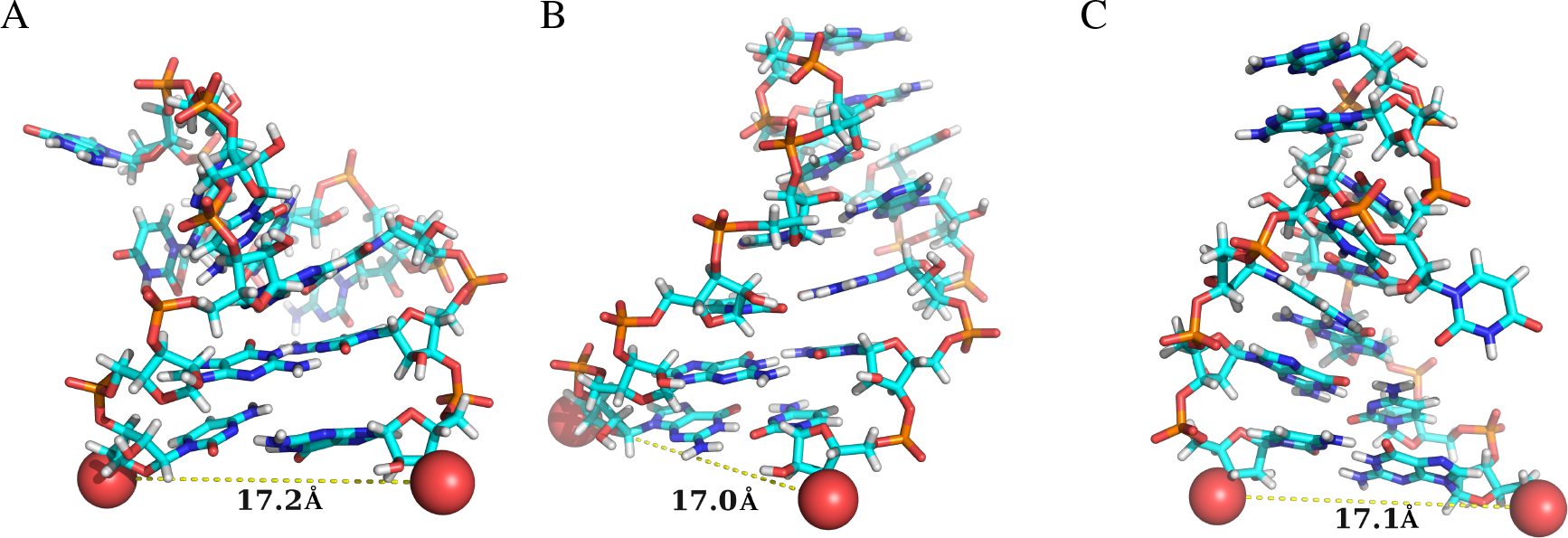
The starting structures for the pulling denaturation simulations. Panel A depicts sequence 1, with loop sequence CAGUGC. Panel B depicts sequence 2, with loop sequence GUAAUA. Panel C depicts sequence 3, with loop sequence UUAAUU. Structures are shown as sticks with carbons in cyan, nitrogens in blue, hydrogens in white, oxygens in red, and phosphoruses in orange. The oxygens used to measure the end to end distance are shown as spheres, and their native distances are indicated with dashed lines and by the boldface number in Å.

We used three hexamer RNA hairpin loops for which NMR structures and optical melting data were both available. For each NMR structure, we used the first conformer provided in the PDB. Below we provide the sequence, where underlining is used to indicate the residues in the loop, as well as references to the deposited coordinates and to the reported stability of each loop.

1. CGA <u>CAGUGC</u> UCG, corresponding to the loop structure with PDB ID of 1AQO (Fig. 1a).^59^ We excised the hairpin loop and closing base pair from the larger structure (CAA <u>CAGUGC</u> UUG, residues 10-21). To stabilize the stem we changed the identity of the nucleotides in the closing helix to two GC pairs by manually changing atom types. This hexaloop caps the iron responsive element that regulates translation of iron associated transcripts by changing their availability to the translation machinery. The experimental free energy change of unfolding of the adjusted structure at 37°C is −0.21 ± 0.39 kcal/mol.^60–62^
2. GGC <u>GUAAUA</u> GCC. We used the first conformer from a solution NMR structure reported by Fountain et al. (Fig. 1B).^63^ The occurrences of this loop in rRNA tend to have a sugar-Hoogstein GA pair between the first and last residues of the loop,^64^ however in the NMR structure this non-canonical interaction is only partially present.^63^ The experimental free energy change of unfolding at 37°C is −3.46 ± 0.24 kcal/mol.^65^
3. GCG <u>UUAAUU</u> CGC, corresponding to the solution NMR structure deposited in the PDB as 1HS3.^66^ The experimental free energy change of unfolding at 37°C is −2.36 ± 0.32 kcal/mol.^67^

We will refer to these chemical systems by the sequence of their loops (for example, sequence 1 will be called CAGUGC).

There are two approaches to estimate experimental uncertainty in folding free energy from optical melts:

1. The average uncertainties observed for the ΔH° (12%) and ΔS° (13.5%) for the database of melts following equation 11 of Xia et al.,^68^ which accounts for the correlation in ΔH° and ΔS°.^62,68^
2. The uncertainty in the free energy change provided with the melt.

To be conservative, we calculated both uncertainties for each sequence and took whichever was greater. We used the statistical method for CAGUGC and UUAAUU, and the uncertainty provided with the melt for GUAAUA.

Sequence 1 was edited as compared to the full NMR structure. Nearest neighbor rules were used to estimate the free energy of unfolding for the revised sequence by subtracting the nearest neighbor terms from the melted sequence’s free energy, then adding the nearest neighbor terms associated with the altered sequence.^68^ Because these modifications were made in an A-form helix, where the nearest neighbor rules are most accurate, the error from the nearest neighbor parameter terms themselves was 0.12 kcal/mol, which resulted in a 0.02 kcal/mol increase in the propagated error estimate of the optical melting experiment.^67^ We used the error estimates, including parameter covariance, to estimate the overall error.^69^

### Simulation Protocol

We chose to monitor and bias folding along the distance between the 5′ and 3′ hydroxyl oxygens of the terminal residues of each 12-mer. This simulates the effect of applying force to the hairpin through tethers attached to its 5′ and 3′ ends, as has been done in single-molecule experiments using optical tweezers.^58^ Here we simulate this transition with umbrella sampling, whereby the molecule is restrained to occupy specific positions along the reaction coordinate in independent equilibrium simulations called windows.^26,70,71^ The biased distributions of reaction coordinate positions are then unbiased to determine the Free Energy Curve (FEC) using the Weighted Histogram Analysis Method (WHAM) to iteratively minimize the statistical errors in overlapping regions from each simulation.^72^

Simulations were run using the GPU code^73^ from the AMBER 14 and AMBER 16 packages.^74,75^ We simulated four replicas of each window for each molecule. Equilibrium distances were spaced in 1 Å increments from 15 Å to 60 Å, and a harmonic restraint with a force constant of 3 kcal mol^−1^ Å^−2^ was used to bias sampling at each distance. The native distance occupied by the NMR structures chosen was approximately 17 Å for each structure (Fig. 1). The first window was taken directly from the solution structure. Subsequent windows were generated by solvating the molecule’s structure from a previously built window, using the last frame of 2 ns of fully solvated MD in the previous window. This procedure was repeated independently for each replicate, so that the starting structure for each replica of a given window is different.

Each window was built with truncated octahedral periodic boundary conditions with a 10 Å minimum distance between the solute and the periodic boundary. Each window was solvated with TIP3P water with neutralizing sodium and with an additional 1 M NaCl to simulate the conditions of the melting experiments. Each system was maintained at 310 K using Langevin Dynamics with a timestep of 2 fs and with bonds involving hydrogen constrained by SHAKE.^76,77^ The collision frequency of the thermostat was 2 ps^−1^ and pressure was maintained using a Monte Carlo barostat with a 100 step attempt frequency. Each simulation was run for 1 μs. Because there were 46 windows per replica of each pulling simulation, and there were four replicas per molecule, this amounts to 4 × 46 × 3 = 552 *μ*s of aggregate simulation time.

### Hydrogen Bond Averaging

To compute the average number of base-base hydrogen bonds we used cpptraj from the Amber-Tools Suite.^74^ We then post-processed the results using awk to exclude backbone and neighboring base interactions. We averaged this quantity across the four replicas for each window, and computed the standard error of its mean (SEM).

### Free Energy Analysis

For each window, the 3′ to 5′ distance was recorded every 10,000 steps (20 ps), leading to 200,000 sampled distances per trajectory. The first 200 ns of each simulation was discarded as equilibration. Umbrella sampling simulations were merged into one free energy curve along the reaction coordinate for each system using WHAM^72,78,79^ with the following parameters: a minimum distance of 14.95 Å, a maximum distance of 60.05 Å, with 451 bins and a convergence criterion for the iterative solution of the WHAM equations of 1.0 × 10^−6^ kcal/mol maximum successive difference for the free energy constants.^79^ The reaction coordinate was broken into two states: slack and stretched (slack includes the native-like windows near 17 Å) divided at 45 Å. We made this choice based on the average number of non-neigboring base base hydrogen bonds (see the section titled Hydrogen Bonding and Reaction Coordinate Partitioning and Fig. 4).

We defined the free energy change of stretching a sequence *A* as:

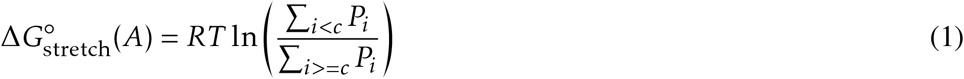

where R is the gas constant, T is the temperature in kelvin *i* is the bin index, *c* is the bin index corresponding to the cutoff distance of 45 Å and *P_i_* is the probability of the i^th^ bin for sequence *A*. To compare the free energy changes estimated from simulations to those measured by optical melting, we used thermodynamic cycles (Fig. 2).^51^ This is necessary because the denatured state is different for pulling than for thermal denaturation.^51,58^ For each pair of free energy changes calculated thus (for sequences A and B), we subtracted the differences from one another to compute:

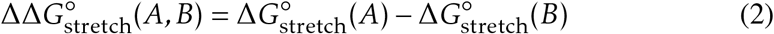

We analogously define a difference in free energy of melting for sequences A and B (where the reference melt for sequence *I* is 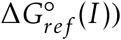:

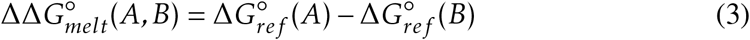

We estimated error using the standard error of the mean of the four replicas.^80^ To compute the variance, the mean, *μ*, was taken to be the 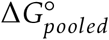 from applying WHAM to the pooled data from the four replicas. We propagated errors using the customary formulas for standard error of the mean (SEM) as the best estimate of the uncertainty in a quantity *q* measured multiple times:^80^

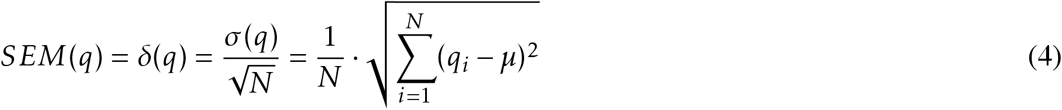

The uncertainty for quantity *Q* calculated by addition or subtraction from other quantities, {*k*}, the values of which are also uncertain, {*δ*_*k*_},is:

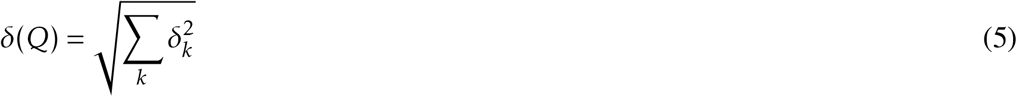

**Figure 2:**
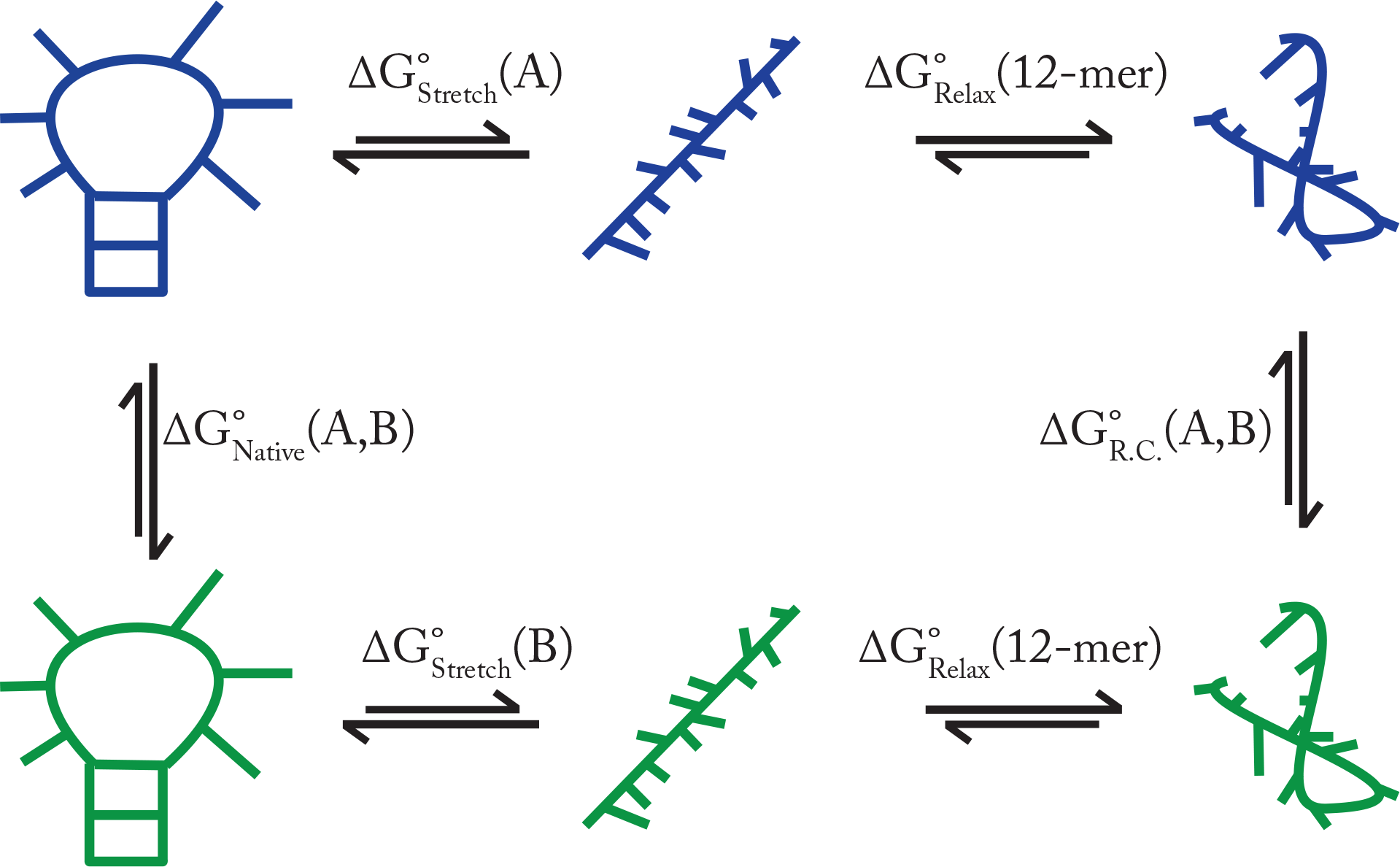
The thermodynamic cycle to compute ΔΔ*G*_stretch_(*A; B*). RNAs A and B. We simulate the transitions from native to stretched, but not the transition from stretched to random coil, the endpoint of optical melting experiments. We can approximate these transitions as sequence independent, here indicated by 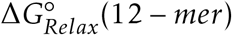, thus they cancel when ΔΔ*G*° are computed.^51^ In the cycle, the free energies are subscripted to show which conformational state the system ends in, when read from right to left. The sequence or sequences involved in the transition is indicated in parentheses after the free energy, thus alchemical transitions are given as involving both sequences. R.C. stands for Random Coil.

Thus the uncertainty in the ΔΔ*G*° described above and characterized by *N*_*A*_ independent simula-tions of Δ*G*° (*A*) and *N*_*B*_ independent simulations of Δ*G*°(*B*) is given by:

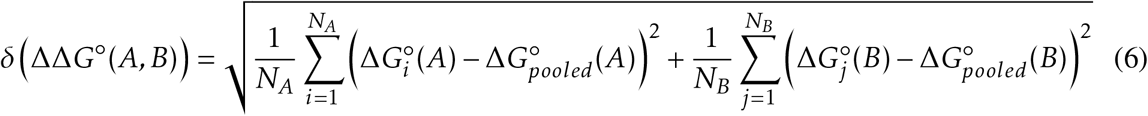

### Assessment of Convergence

#### Trajectory Truncation

To estimate the amount necessary to exclude from sampling as equilibration, we truncated the trajectories with increasing portions of each window trajectory from its beginning. As this equilibration is extended, the sampling is more distant from the system-building procedure and thus less perturbed by it. We truncated between 0 and 600 ns from each trajectory, in steps of 100 ns. We then used these FECs to calculate a series of 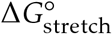.

In a second analysis, the trajectories were truncated from the end, again in 100 ns increments and removing between 0 and 600 ns. This analysis exposes how our results would vary if we had stopped the simulations earlier.

#### Covariance Overlap by Window

We used a singular value decomposition (SVD) to solve for the the eigenvalues and eigenvectors of the atomic motions in the simulations representing each window, then computed the covariance overlap of the eigenvectors to evaluate their similarity:^81–83^

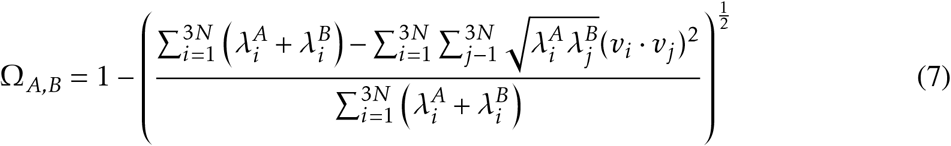

where Ω_*A,B*_ is the covariance overlap between trajectories *A* and *B* for *N* selected atoms, 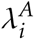 is the *i^th^* eigenvalue of the covariance matrix from *A*, and *v_i_* is the corresponding *i^th^* eigenvector. We used algorithms to perform the SVD and the covariance overlap in the Lightweight Object-Oriented Structure analysis library project, LOOS.^83,84^

For each window, all heavy atoms in all four trajectories were aligned consistently.^82^ SVDs were then computed for each replica of each window. The covariance overlap was then computed for each non-redundant pairwise combination of the four replicas. We averaged the resultant per-window scores and estimated the quality of that average by computing an SEM where the number of replicas (not the number of combinations of replicas) determined the sample size.

#### All-to-all RMSD of Neighboring Windows

For WHAM, sequential windows must contain overlapping probability distributions along the reaction coordinate, otherwise WHAM will produce a discontinuous free energy curve. The theoretical basis of umbrella sampling, however, is repeated free energy perturbation, so the windows must overlap in a more complete phase space.^70^ Overlap along our reaction coordinate is good as evidenced by our smooth FECs. To score phase-space overlap in dimensions not well characterized by our reaction coordinate, we assessed the similarity of loop structure in consecutive windows by computing a root-mean-squared displacement between each frame (an ‘all-to-all’ RMSD) in trajectories from neighboring windows for all heavy atoms in the loop (residues 3-9). We wrote a LOOS tool called trans-rmsd to perform the calculations between frames from two user specified sets of trajectories, and have included it with the prebuilt tools that package ships with.^84,85^

To score similarity between windows, we consider a fraction of frame pairs whose RMSD is below a chosen cutoff (2.5 Å). RMSD between loop heavy atoms below this cutoff implies similar conformations. This quantity could be computed for any two windows, or indeed any pair of trajectories. We refer to the fraction, *F*, for two windows (named by their window indices) *v* and *w*, as *F*(*v, w*). Because we are interested in scoring overlap along the reaction coordinate, we only consider neighboring windows, *F*(*w, w* + 1). We also computed a ratio of the neighboring fraction of frame pairs below the cutoff to the average fraction of frame pairs within the two neighboring windows that are below the cutoff:

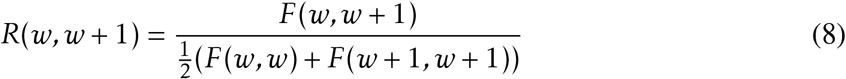

This ratio is a clearer representation of the degree of overlap where *F*(*w, w* + 1) is small (less than 0.1), because *F*(*w, w*) and *F*(*w* + 1, *w* + 1) are also small. This occurs when windows with high conformational heterogeneity are compared. We used pooled frames from all four trajectories of each window sampled every 500 ps to compute these values.

### FEC Numerical Derivatives

To evaluate which bins along the reaction coordinate exhibit greater variation in free energy, we computed a numerical derivative of the FEC as the difference between neighboring bin free energies (where the bin with lower index is subtracted from the bin with higher index) divided by the bin width. We then computed the standard deviation of this quantity, σ, across the four replica distributions of each bin, where we took the mean to be the free energy resultant from the pooled distributions.

For any FEC, *G*, with distance between points δ, and free energy at the *i^th^* point 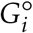:

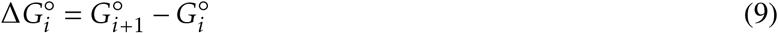

Then the numerical derivative between these points is:

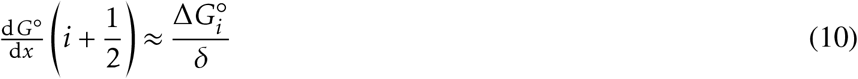

where the argument indicates that the expression approximates the derivative between points *i* and *i* + 1. We now have a force along the FEC, and can consider the spread in force per point between runs by calculating the standard deviation in this quantity across *N* replicas for all the neighboring pairs along our reaction coordinate, where 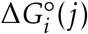 represents the jth replica. 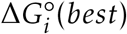 is the value of the neighboring point difference computed with the pooled distributions.

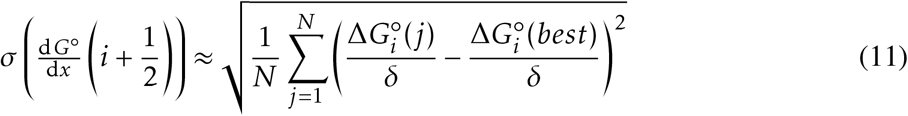

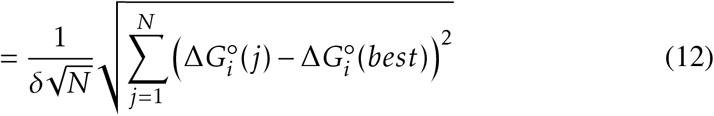

To quantify these values per window, we collected the FEC derivative curves into segments where the center of the segment was the restrained distance of the associated window. We assumed each segment should be 1 Å wide, in keeping with our restraint spacing. We then integrated over these segments using the trapezoid rule as implemented in NumPy.^86^ This quantity is in units of energy, and computing its standard deviation provides a score of the variability observed within a window. We provide the Python script we used to perform these calculations on FEC data files as Supplementary Materials. This method is general to any comparison between curves with arbitrary zeros, where the same problem (of offsets in parts of the curve proximal to the zero rendering a variance distal to the zero incoherent) would apply for repeated measurements of the same curve just as it does here.

## Results

### Free Energy Curves

We obtained free energy curves for the three sequences (Fig. 1) using WHAM as applied to our umbrella sampling simulations (Fig. 3, panels A, C, and E). Our best estimate for the Free Energy Curves (FECs) exclude the first 200 ns of sampling for equilibration and pool the sampling from each replicate of each window before running WHAM. This best estimate is shown in black, and is considered the full length dataset. Because the simulations were done in quadruplicate, overlaying the FECs shows regions that disagree across replicas. While agreement does not guarantee convergence, disagreement is necessarily the result of insufficient sampling, though not always in the same bin as the disagreement (see the section on *Per-Bin Free Energy Variation* below). Thus it appears that our simulations provide consistent representations of the native basin, the region surrounding the native structure at approximately 17 Å, and of the extended structures present beyond 45 Å. This is unsurprising; these regions of the reaction coordinate should correspond to relatively small volumes in phase space by comparison with the middle of the reaction coordinate, where the native stem is lost but the backbone is not taut.

To assess convergence, we removed increasingly large sections from the end of the trajectory of each replica in 100 ns increments before pooling them and applying WHAM, as before (Fig. 3, panels B, D and F). This addresses the extent to which the ‘best estimate’ provided by the pooled data is sensitive to the length of the simulations. We note that these curves exhibit some disagreement when they are zeroed at their minima, which again is at 17 Å. We quantify the extent to which this disagreement affects the calculation of Δ*G*° values from these data in the subsection on *Free Energy Changes* below.

### Hydrogen Bonding and Reaction Coordinate Partitioning

To identify the endpoint of stretching, we computed the average occupancy of hydrogen bonds between non-neighboring bases along the reaction coordinate. We found that this average consistently drops to 0.3 hydrogen bonds or below at 45 Å and, with the exception of the 53 Å window from the first replica of CAGUGC, remains near zero along the rest of the reaction coordinate (Fig. 4). By monitoring all hydrogen bonds we considered non-native structures forming in windows that are near the native basin, but distant enough to have disrupted the helix, as non-stretched structures. This can be seen from the heterogeneity in the hydrogen bond plots for each sequence (there is also variability amongst replicas). The most consistent pattern is the near absence of such hydrogen bonds beyond 45 Å, and we therefore chose this as the boundary between slack and stretched. While the boundary would move for hairpin sequences of different lengths, this heuristic could still be used to identify a boundary for such systems.

**Figure 3:**
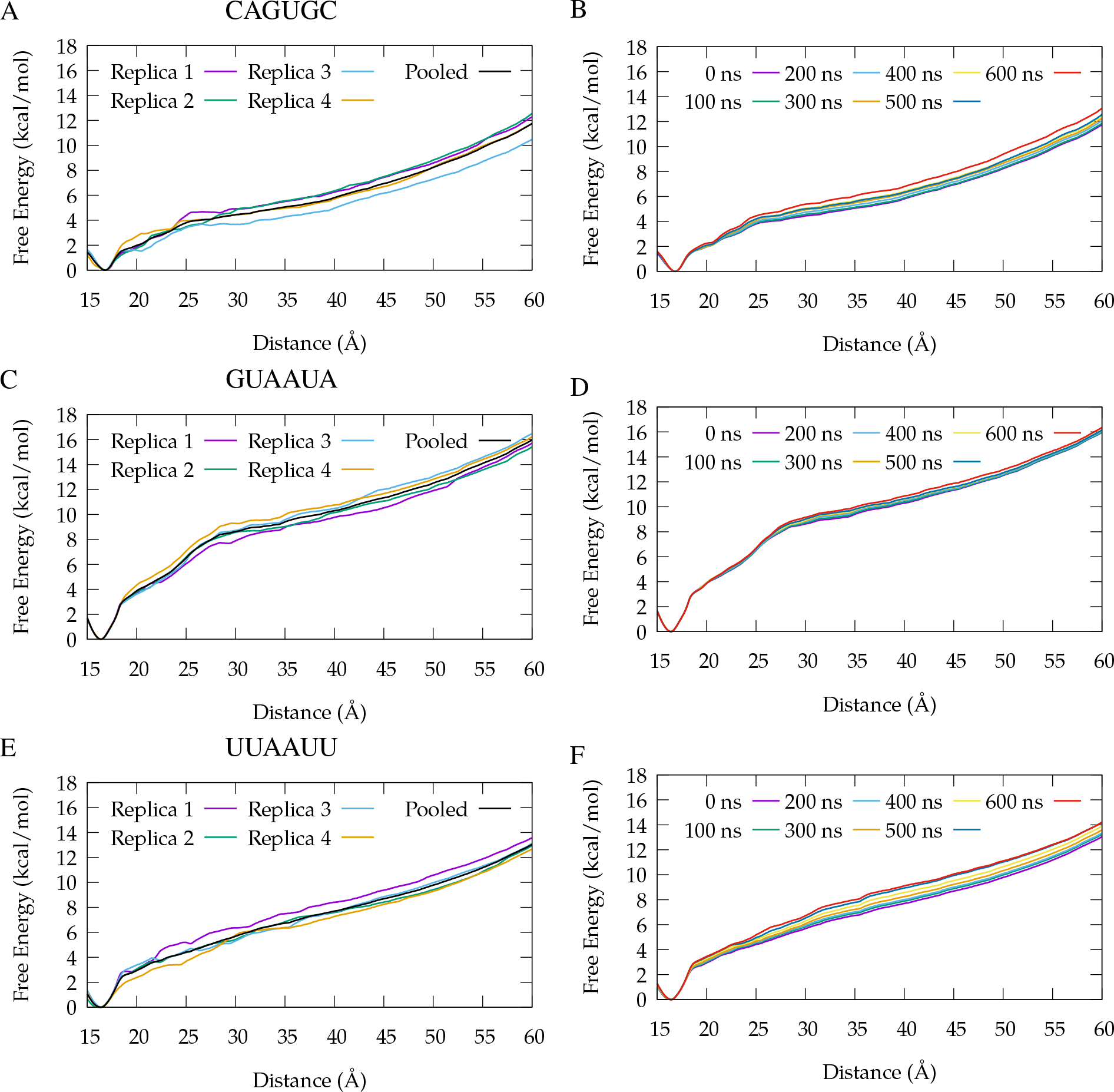
Free Energy Curves. Panels A, C, and E show the four replicas of the equilibrium-excluded data for each system, plus the FEC produced by pooling that data, which is our best estimate for the FEC. Panels B, D, and F show the results the pooled data from trajectories with data excluded from their ends in 100 ns increments. Panels A and B correspond to CAGUGC, C and D correspond to GUAAUA, and E and F correspond to UUAAUU. For each truncated FEC, the curve corresponding to ‘0 ns excluded’ is the same as the pooled data in the plots showing the overlay of the replicas of each system. Note that all FECs are zeroed at their minimum value, which happens to be the same (the location of the native state, 17 Å) for all 12 FECs.

**Figure 4:**
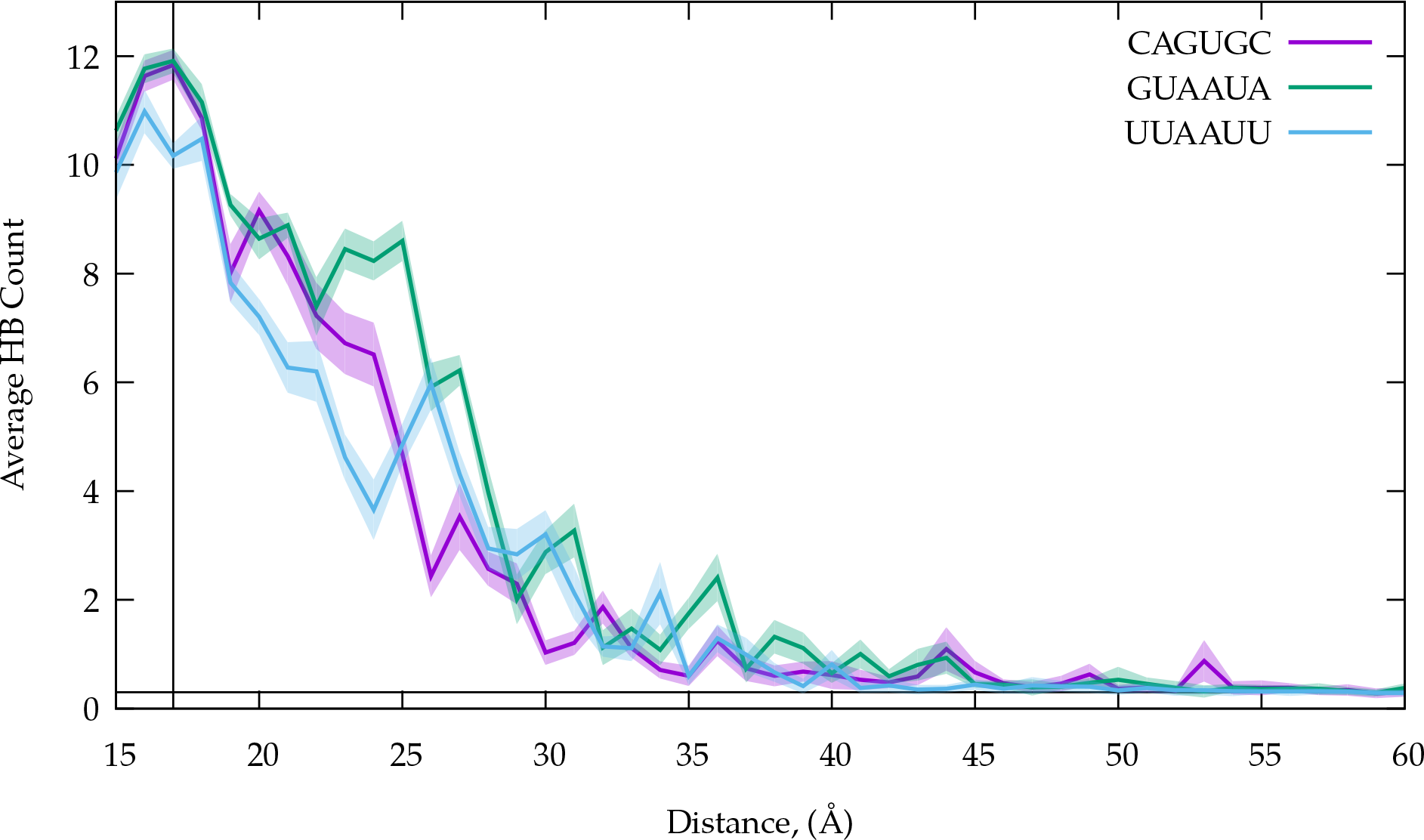
The average hydrogen bond occupancy per window for hydrogen bonds between donors and acceptors on non-neighboring bases. The transparent envelopes represent the standard error of the mean (SEM). The hydrogen bonds were defined by an angle cutoff of 30° and a heavy atom distance cutoff of 3.5 Å. The horizontal line is at 0.3 average hydrogen bonds. The vertical line at 17 Å indicates native Watson-Crick helix.

### Convergence and Overlap of Windows

Figure 5 shows the controls we performed to further address convergence and window overlap. We computed the covariance overlap (Eq. 7) for each replica of each window, and report an average of these with an SEM as an envelope around each point in Panel A. The covariance overlap gives a normalized distance between two covariance matrices (it is undefined for non-analogous selections of atoms). A covariance overlap of 1 would indicate that the covariance matrices are identical, whereas a covariance overlap of 0 would indicate that there is no similarity between the two covariance matrices.^81^ Because the key component of this distance is defined as the dot product of all pairs of eigenvectors from the covariance matrices for the two simulations (the principal components of the motions) these scores can also be viewed as measuring the extent to which the principal components overlap.^83,87^ For perspective, we computed the covariance overlap for just an alignment and PCA based on the helical residues of the native window. We view this as a best-case covariance overlap for a structured RNA because the ends of the helix are restrained to be at the native distance and the helix should be less flexible than the loop, although the loop-closing bases and the helix closing bases can still fray. The average and standard deviation of these calculations was 0.55 ± 0.19. We conclude from this that there is still heterogeneity in the loop motions in restraint distances shorter than 45 Å. However, by comparison with our helical reference few of our windows exhibit disparate motions, and our more extended windows exhibit remarkably similar ones. We also calculated the subspace overlap of the first 25 modes of the neighboring simulations (see Supplementary Fig. 1); they were uniformly higher than the covariance overlaps, which is not surprising because the subspace overlap treats the eigenvectors as unit vectors.

**Figure 5:**
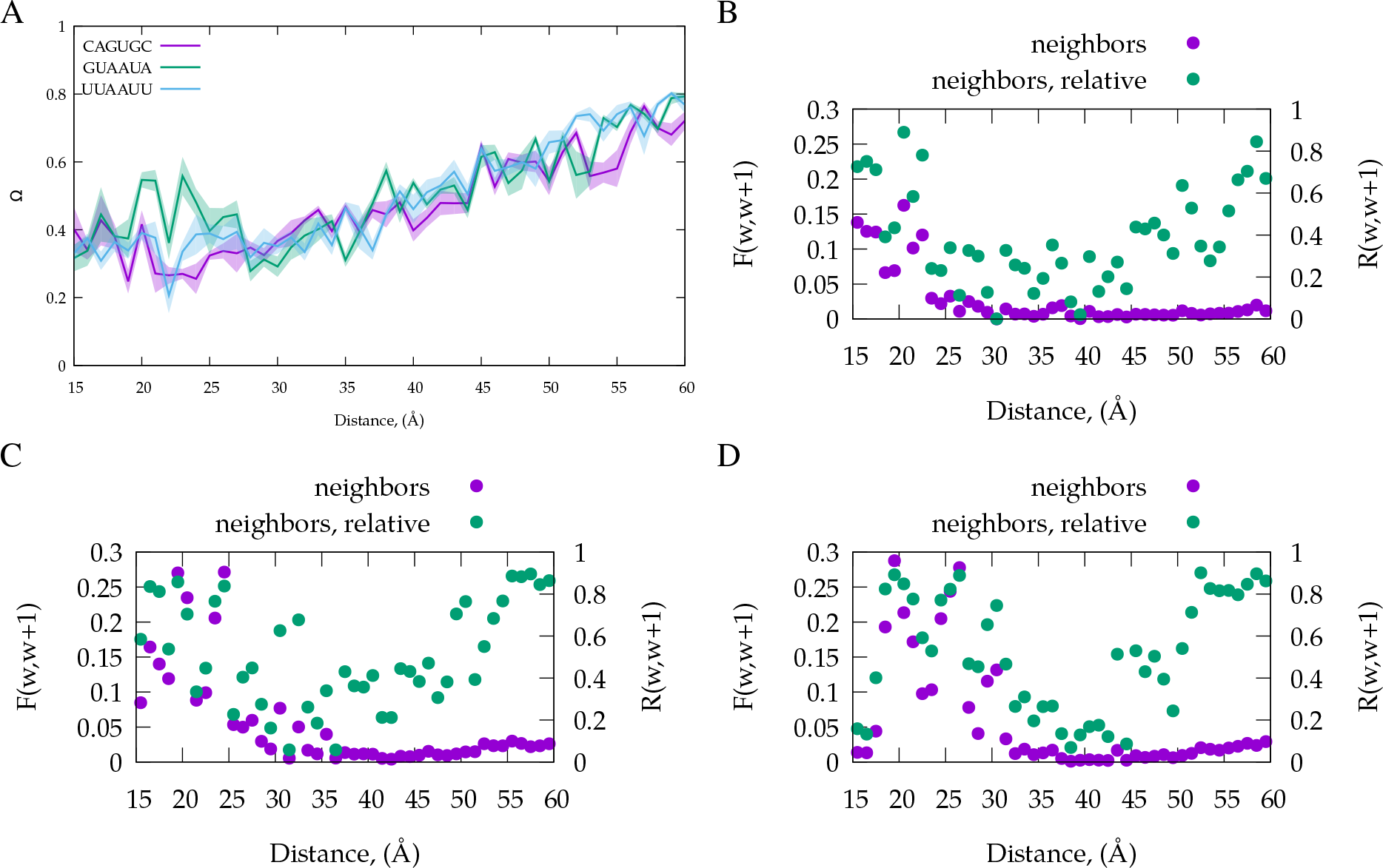
Assessment of convergence and phase space overlap. Panel A shows the covariance overlap between the pooled trajectories for all heavy atoms in the loop. The bold line indicates the average value, with the transparent envelope indicating the SEM Panels B, C and D, corresponding to CAGUGC, GUAAUA, and UUAAUU, respectively, plot the fraction of pairs of frames with a pairwise all-heavy-atom loop RMSD less than 2.5 Å that occurs between each window *w*, and its neighbor *w* + 1, *F*(*w, w* + 1) is show on the left axis. The relative fraction, *R*(*w, w* + 1) = 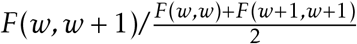, is shown on the right axis.

To assess phase space overlap between windows, we computed the fraction of all frame pairs in neighboring windows with loop heavy atom RMSD below 2.5 Å (Fig. 5, panels B, C, and D). All three molecules have substantial fractions of such frames from neighboring windows before 25 Å. After 30 Å, the fraction drops below 0.05. Beyond this point there is no native structure, and most conformations are only observed for a single frame. To calibrate this fraction we also computed a ratio between the fractions of frames below the cutoff from neighboring windows and that same fraction from the average of both neighboring windows compared to themselves. It scores overlap relative to how heterogeneous the window under consideration is, which is useful in understanding the region of low overlap in the middle of the reaction coordinate. These quantities are computed between neighboring windows. For each window, the trajectory analyzed was the pooled frames from all four of the replicas, sampled every 500 ps. Interestingly, while both scores are below 0.01 for the middle windows, for the higher windows near the ‘stretched’ state they begin to rise again, suggesting those windows have more similar conformational ensembles.

### Per-Bin Free Energy Variation

In the free energy curves in Figure 3, we noted a pattern in the disagreements between curves from the same sequence; disagreement usually occurs starting at a certain position on the curve, the distance between a truncation and its neighbors (or a replica and its neighbors) rises, then begins to mirror its neighbors again. Because variability at a certain bin produces an offset in bins more distant from where curves are zeroed across replicas, we consider a numerical derivative that will only score variability between two neighboring bins (see the Methods subsection *FEC Numerical Derivatives*).

In Figure 6, the regions of lowest variability are near the native distance where the replicas are overlapped in Figure 3. There is a region of low variability in the windows that also lack non-neighboring base-base hydrogen bonds (Fig. 4). Bins from the ‘stretched’ state tend to be less variable than those in the extended but not taut middle region of the curve. But note the variability in CAGUGC that corresponds to a rearrangement in the base pairing of the terminal base pair in shorter than native distance (compressed) windows in some replicas

The window integrals of the standard deviation (Fig. 6, panels B, C and D) suggest the extent of convergence for individual windows with respect to their part of the FEC. Low variability windows have almost no (less than 0.1 kcal/mol) difference between the truncated and full trajectories, and the total variability estimated from truncations also remains low (below 0.15 kcal/mol). Windows that vary substantially can be twice as variable for truncated trajectories, and even with the full dataset score higher than 0.2 kcal/mol.

### Free Energy Changes

To estimate how much of the trajectory to exclude as equilibration, we plotted the free energy of stretching (Eq. 1) from end-distance distributions that had been truncated from the beginning in 100 ns increments in the panel A of Figure 7. While it is challenging to determine how much equilibration is sufficient and not excessive, for two of our three systems it appears that the free energy of stretching stops systematically changing after the first 200 ns. For UUAAUU, there appears to be a steadier drift in the value of free energy that occurs over the course of the trajectories, although it leveled off considerably after excluding 200 ns. Based on this evidence, and by inspecting our FEC derivatives as a function of truncation from the beginning, we conclude that a 200 ns equilibration is sufficient.

**Figure 6:**
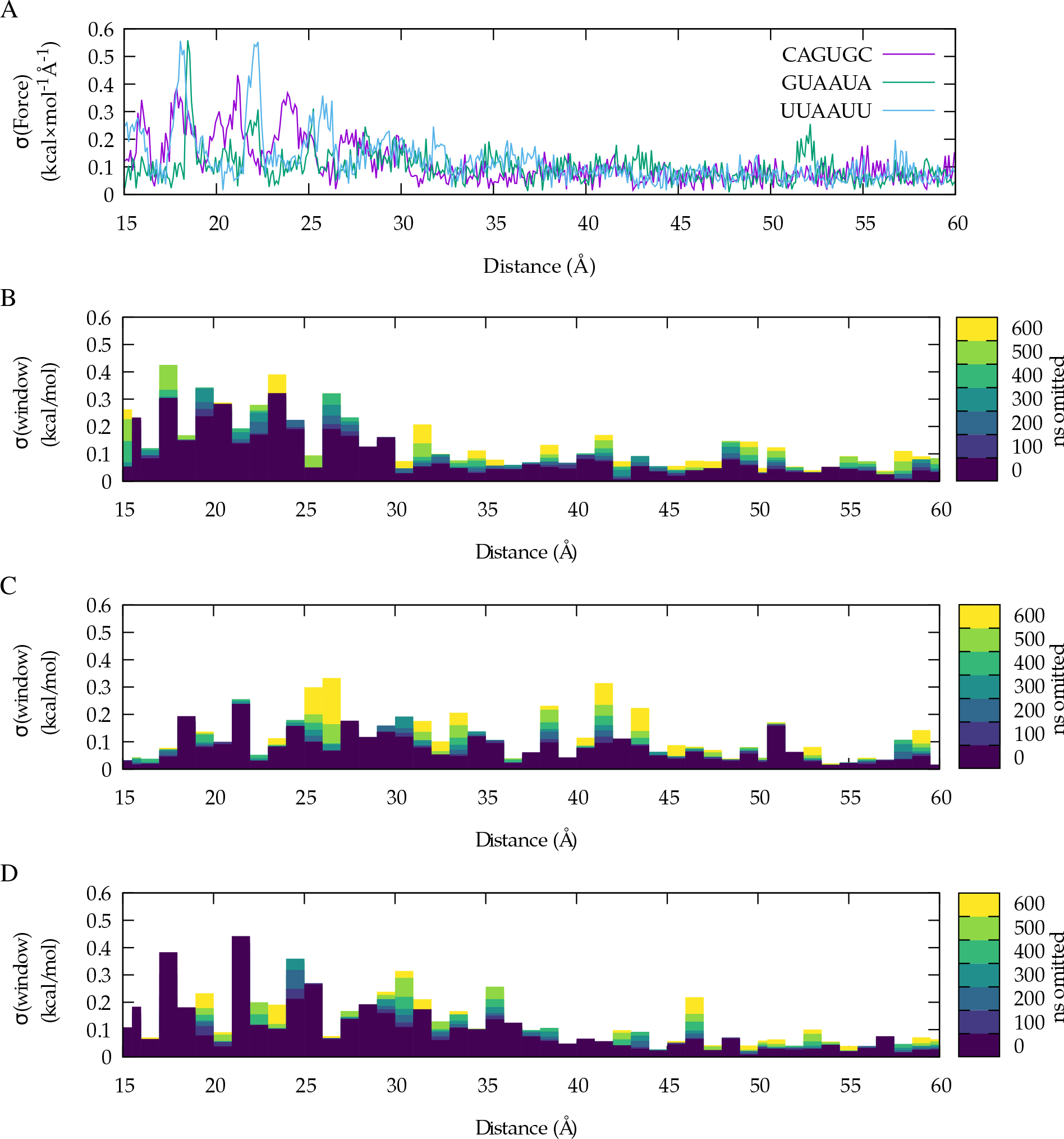
The standard deviation σ per bin in the FECs. Panel A shows σ for the derivative of each FEC with respect to the reaction coordinate (see Methods subsection *FEC Numerical Derivatives*) for data derived from the full length trajectories. In panels B-D, the regions corresponding to each window are integrated along the reaction coordinate using the trapezoid rule, and plotted as boxes. Panel B corresponds to CAGUGC, C to GUAAUA, and D to UUAAUU. The longer trajectories (fewer ns omitted from the end) are overlaid on the shorter trajectories.

**Figure 7:**
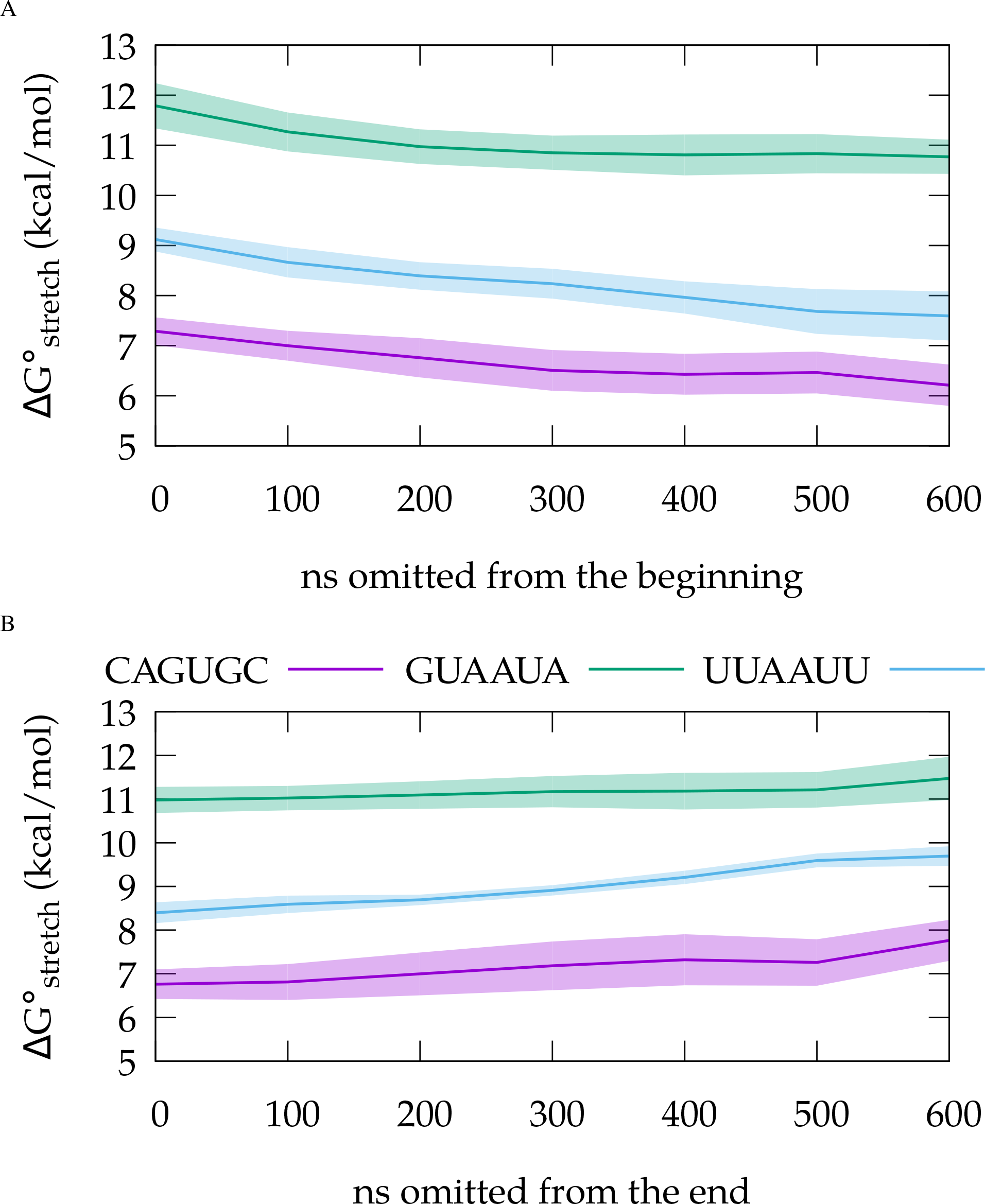
The 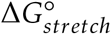 as a function of truncation. The magenta line and envelope correspond to CAGUGC, the blue to UUAAUU, and the green to GUAAUA. The lines correspond to the FEC resulting from the pooled data truncated by the amount on the independent axis. The envelope is the SEM of the pooled data with the truncated replicas treated as independent samples. Panel A shows the relaxation of the 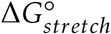 from the complete dataset truncated from the beginning. Panel B shows how the same quantity varies as a function of data excluded from the end of the full-length (equilibration excluded) data.

**Table 1:**
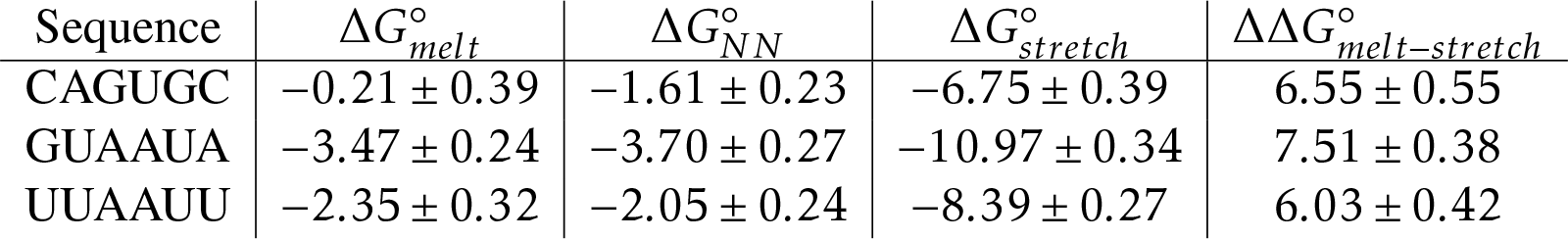
The free energy changes (kcal/mol), measured, predicted using the nearest neighbor (NN) parameters,^61,69^ and simulated, for each sequence with uncertainty for each value. The 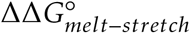 quantifies the error in the thermodynamic cycle calculations because that quantity should be a constant up to the variability from incomplete sampling. The mean with propagated error for this quantity is 6:70 ± 0:79 kcal/mol.

To compare our stretching experiments to experimental folding stabilities (Tables 1 and 2), we followed our approach from Spasic et al.,^51^ where a thermodynamic cycle describing differences in folding stability (Fig. 2) was used. By approximating the stretched to random-coil transition as sequence independent, as is done in single-molecule pulling experiments,^58^ we can compare the ΔΔ*G*° between two sequences to analogous differences in stability from optical melting experiments. To quantify the sequence independence of the stretched state from our data, we fit the free energy curves to quadratic functions in the stretched region (45 to 60 Å), and found that the quadratic and linear coefficients are similar (Supporting Table 2). This lack of variability in the stretched region of the curve supports our approximating it as sequence independent; if the curve were variable in this region it would be harder to distinguish whether this variability were a property of the sequence or whether it was random variability from sampling.

Table 1 shows the changes in Δ*G*° we observed. As discussed in the Introduction, the changes in state between the *in silico* stretching and *in vitro* melting are not the same, so it is expected that these free energies would be different. Our assumption that the differences in stabilities between sequences (Table 2) should be comparable whether the stabilities were obtained from melts or stretches also implies that there is some approximately constant component of the stretching free energy that is a sequence-independent offset. When we compute relative stabilities using our thermodynamic cycle, the sequence-independent components cancel. In the column labeled 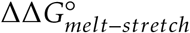 we estimate this quantity by taking the difference between the 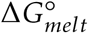 and the 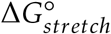 for the same sequence. In principle, if there were no sequence dependence and no sampling error, these numbers would be the same for all three sequences. We assessed statistical errors in our study using technical replicas, therefore differences in this quantity that are outside our uncertainties can be interpreted as systematic differences. We estimate the difference (Table 2; Melt-Stretch) to be similar for UUAAUU and CAGUGC, but that for GUAAUA appears to be the outlier. It is interesting that the similar comparison between stabilities estimated using the nearest neighbor model^88^ and those obtained from optical melting experiments (Table 2; Stretch-NN), the largest outlier is CAGUGC. Our simulations appear to model the instability of this loop although the nearest neighbor model does not.

**Figure 8:**
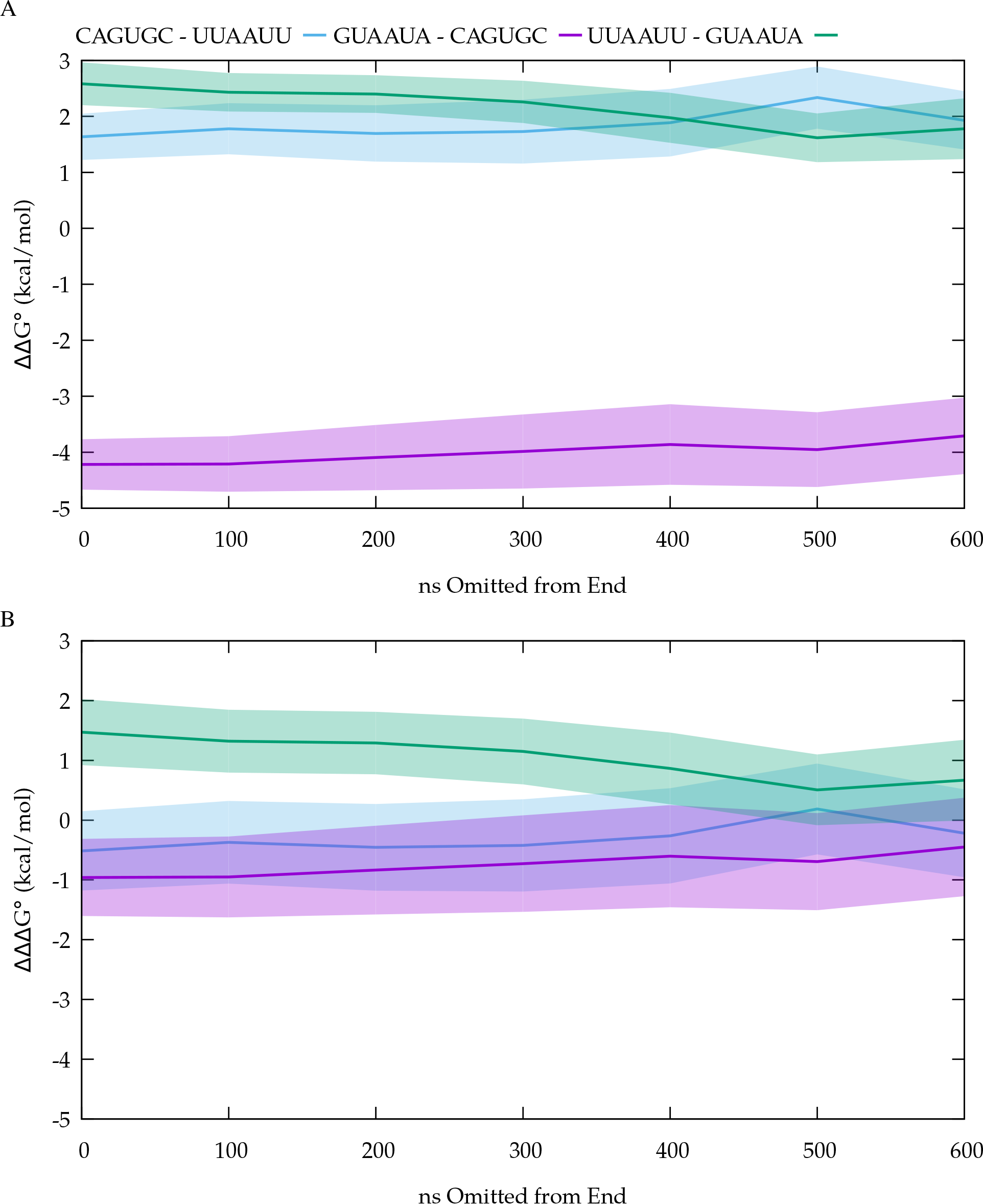
The 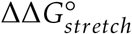 and residuals as a function of truncation. Panel A shows the 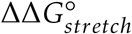 with SEM from replicas. Panel B shows the residual between that value and its reference ΔΔ*G*° with propagated error.

Because the residuals average to 1 kcal/mol and the errors for individual residuals are of order *k_B_T*, we conclude that we are nearing chemical accuracy and precision, respectively, with this technique. We propagated uncertainty in both the physical measurements and the experiments to a residual to determine whether these values are significantly different. If they are not, the residuals 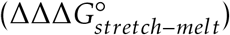) would be close to zero in free energy, and zero would be within the error-envelope of the values. Based on Table 2 and Fig. 8, we cannot reject the null hypothesis that the residuals are zero for two of the three residuals. This is also true even in highly truncated trajectories (Fig. 8), and this indicates less sampling would have reached the same conclusion. The deviation from zero residual seems to vary by sequence—the two residuals involving GUAAUA are more erroneous. While this can likely be explained in part as systematic (rather than sampling) error in these free energy differences, the differences from zero are not significant (Table 2).

**Table 2:** The difference in free energy change (kcal/mol) of unfolding for pairs of sequences. Each of these differences are from the thermodynamic cycle in Fig. 2. The differences are calculated as alchemical transitions between sequences, with 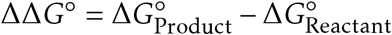. The ‘stretch’ transitions are those calculated from simulations, and the melting transitions are our reference data from the literature. 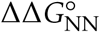 is what the nearest-neighbor model estimates for each sequence pair. The column Melt-Stretch is the difference between the experiment and the simulation, referred to as the residual. Stretch-NN is the difference between 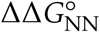 and 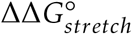. The uncertainties given for the residuals are the uncertainties propagated from the ΔΔ*G*° using Eq. 5. The P values are provided in parentheses after the residuals and are the probability of the null hypothesis that the differences are zero given the data. They are computed from a two-tailed t-test against a particular value, zero, with three degrees of freedom.

| Reactant | Product | $\Delta\Delta G^\circ_{\text{stretch}}$ | $\Delta\Delta G^\circ_{\text{melt}}$ | $\Delta\Delta G^\circ_{\text{NN}}$ | Melt-Stretch (P) | Stretch-NN (P) |
| --- | --- | --- | --- | --- | --- | --- |
| GUAAUA | UUAAUU | $2.58 \pm 0.44$ | $1.11 \pm 0.40$ | $1.65 \pm 0.35$ | $1.47 \pm 0.60(0.90)$ | $0.93 \pm 0.57(0.198)$ |
| UUAAUU | CAGUGC | $1.64 \pm 0.48$ | $2.15 \pm 0.52$ | $0.44 \pm 0.33$ | $-0.52 \pm 0.70(0.514)$ | $1.20 \pm 0.58(0.131)$ |
| CAGUGC | GUAAUA | $-4.22 \pm 0.52$ | $-3.26 \pm 0.46$ | $-2.09 \pm 0.36$ | $-0.95 \pm 0.70(0.264)$ | $-2.13 \pm 0.63(0.044)$ |

## Discussion

### FF99+bsc0+χ_OL3_ Brackets the Relative Stability of RNA Hairpins

This study was designed to determine whether there is systematic error in the free energies simulated for RNA hairpin pulling. There is a growing consensus that the RNA force fields for standard molecular dynamics are inadequate, but this is commonly evaluated in a structural context with the observation that simulations do not preserve native conformations.^24,32^ The argument could be made that folding stability could be approximately correct even if the balance of conformations were incorrect, if the number of microstates available to the RNA and their relative energies were approximately correct. Our results suggest that FF99+bsc0+χ_OL3_ can perform with chemical accuracy to estimate relative stabilities of RNA hairpins along an end to end reaction coordinate with adequate sampling. While convergence is never promised when sampling on a rough free energy surface, these results suggest that in spite of the parameter imbalances discussed in the literature, the folding free energies of RNA hexaloops can be adequately modeled.^24,32,33,37^

This work also provides a framework for benchmarking force fields that displays incremental progress, which is a current challenge. Consider for example comparisons between simulations and two dimensional Nuclear Overhauser Effect (NOE) spectra.^24^ It is clear that a simulation indicating the presence of strong NOE cross-peaks that are not observed in experimental spectra of the same system is a problem. What is not clear is how inaccurate this comparison shows the force field to be. It is conceivable that minor adjustments in the right parameters would tune the force fields and eliminate these artifacts entirely, although we find this unlikely given the breadth of efforts, including our own, to rectify these issues.^40^ Such efforts have in some cases resulted in force fields that sample structures with unobserved NOEs less frequently, but do not totally reduce them to levels that would be undetectable by NMR. If the difference in free energy between a minor conformation and a major one is mis-estimated by the force field by as little as 2.0 kcal/mol at 277 K, it could be the dominant conformation in the simulation and so rare as to be unmeasurable by NMR. Because we cannot measure the features of the free energy surface of the tetramer systems experimentally, it is challenging to differentiate between force fields that are becoming more accurate from those that are introducing new errors. In particular, it is hard to determine whether a given modification is part of a necessary set of changes to the force field, but not in its own right sufficient. The progressive nature of free energy benchmarks therefore makes them a needed addition to the nucleic acid force field test suite.

### Convergence

In principle, molecular dynamics simulation (MD) allows for the sampling of the thermodynamic ensemble of the simulated system. This ensemble could then determine any equilibrium quantity of interest and be compared with or estimate experimental data. In practice, especially because of technological limits, sampling is usually incomplete, causing statistical errors in results.^23^ For RNA simulations, the statistical problem is pronounced; while protein folding simulations have been the focus of pioneering work in enhanced sampling,^89^ swarms-of-trajectories approaches that harness the aggregate idle power of many personal computers,^90^ and most recently brute force sampling by specialized supercomputers,^91^ similar simulations of RNA have either only been tried recently,^32^ or have not yet been performed. Because of this lack of focus by the broader MD field, the source of reported inaccuracies is not obvious.

The RNA folding landscape is not completely defined by the end to end distance. For specific windows in umbrella sampling, there might be slowly equilibrating degrees of freedom that are nearly orthogonal to the reaction coordinate, and some of these degrees of freedom might be poorly sampled. These degrees of freedom must overlap for the results of an umbrella sampling approach to be interpretable, even though the overlap between adjacent windows along the reaction coordinate is specifically what determines whether WHAM produces a continuous curve. To score the overlap of window ensembles on potentially orthogonal degrees of freedom, we considered all-to-all frame RMSDs for nearest neighbor and next-neighbor windows along the reaction coordinate (Fig. 5, panels B, C and D). This RMSD was calculated for all heavy atoms in the loop residues. The loop is the most distal part of the RNA from the restraint, and therefore the most likely to contain degrees of freedom for which the biasing potentials enhance sampling the least.

In the longer distance windows the restraint becomes a more effective bias for all the degrees of freedom. Within more distal windows (those above 25 Å restraint distance) there is still conformational heterogeneity as evidenced by the relatively low fraction of frame pairs below our cutoff, however it appears that these windows exhibit less diverse motion according to the covariance overlap, suggesting that perhaps lever-arm effects from the extended structures are at play in reducing the fraction of frame pairs below the RMSD cutoff.

While the relative fraction of pairwise RMSDs suggests there is less overlap of ensembles in the middle windows, we note that no neighboring windows have zero frame pairs below the 2.5 Å cutoff in RMSD. We also note that this calculation, where only the loop conformations are scored, is a conservative choice. If we had selected all heavy atoms, the molecule overlaps would have been higher because they would have contributions from atoms that are more directly restrained to be in similar positions. Finally, we note that in studying the variation in FEC derivatives per window (Fig. 6), it appears that windows for distances of 26-60 Å (the windows with the greatest conformational heterogeneity by this metric) result in the smallest changes in the free energy curves themselves. Their loop motions are also more self similar, even though the loop structures are not self similar (Fig. 5A). In other words, for the purposes of precisely calculating the free energy of stretching, additional sampling of the middle and terminal windows would yield the least change in the computed free energies. It therefore appears that our reaction coordinate is nearly orthogonal to the loop conformations in the mid-range windows.

One might wonder whether, with more extensive sampling, we might find our residuals trending to zero. While this question is hard to answer, considering the residual as a function of truncation might suggest whether extending our sampling further would result in different conclusions.^92^ From Fig. 8, we see the residuals increase as the trajectory gets shorter, which is what one might expect if instead of excluding unequilibrated parts of the trajectory we were simply excluding viable samples from our dataset.

### The Reaction Coordinate

The major benefit of the end-to-end distance reaction coordinate is the well defined character of the endpoint. Sampling the random coil ensemble for an RNA large enough to have structure is not feasible given the level of access most simulators have to computing power. Thus simulations of stretching transitions are more likely to provide useful estimates of stability given current technology. A minor benefit is that force-driven extension experiments also follow this reaction co-ordinate, although the experiments currently use molecular handles that are are too large to include in our calculations.

Simulations estimating stability from pulling-like restraining forces have historically suffered from the problem of having more arbitrary distinctions between native and stretched states.^38,93^ Using the aggregate non-neighboring base-base hydrogen bonds as a score of extension avoids the problem of searching for a point along the reaction coordinate were native structure has been lost. This point is not always obvious because it is difficult to distinguish between native fluctuations and an unfolded conformation. Our choice is a way to define the point where the chain is stretched, which allows us to lump many of these potentially native-like structures into the slack state. The hydrogen bond metric can be applied to unimolecular stretching experiments of RNAs of arbitrary size and sequence.

### Standard Deviation of Free Energy Derivative

In umbrella sampling, a deviation between two technical replicas is usually particular to specific bins along the reaction coordinate. Given the pattern of variability along the reaction coordinate it might seem natural to consider a standard deviation per-bin. This is problematic because the bins are zeroed at one location along the reaction coordinate. At points along the reaction coordinate distant from the zeroed location, statistical fluctuations will have accumulated. Computing a variance or standard deviation with these values directly would therefore be incorrect.

To assess the agreement in FECs by bin, we computed the standard deviation in the numerical derivative of the FEC across replicas (see section *FEC Numerical Derivatives* in the Methods and Fig. 6). We submit this as the best approach for analyzing FECs, as it provides information about consistency and accurately identifies regions of disagreement across replicas or even across sub-datasets. It would certainly apply to discretized FECs assembled from simulations by other means, such as the Multistate Bennett Acceptance Ratio,^?^ weighted ensemble simulation,^?^ and well-tempered Metadynamics.^?^ Computing numerical integrals of this quantity across the width of a window illustrates how variability is localized to a given window. Both quantities combine nicely with dataset exclusion approaches such as those discussed in the section *Trajectory Truncation*, because they can indicate which parts of a FEC are not as well converged.

Thus, both quantities would find use in further efforts to converge umbrella sampling calculations. The window integral standard deviations indicate windows that need further sampling. To initialize simulations for additional sampling, the derivative can suggest what frames to start from because it is a per-bin score of variability. Frames from a variable window exhibiting distances along the reaction coordinate that would be binned under a peak in the derivative curve are the best frames from which to start new simulations, because they are adjacent to some under-sampled event that has resulted in variability at that position. This could be done to adaptively guide additional sampling while simulations are ongoing.

### Furthering *in Silico* Pulling

A number of approaches could be used to improve the efficiency of these types of *in silico* pulling experiments. Our results specifically suggest:

1. Sample the stretched state less, or not at all. The stretched windows do not contain much information about our systems but are the most computationally expensive because they are larger and therefore have much greater water content. The windows with restraint distances 45 Å and greater are some of the most well converged by our metrics, especially the covariance overlap and the free energy derivative SDs. Because we approximate the stretched state as sequence independent, we could add additional sequences onto our analysis by averaging the stretched windows together and considering only their differences in the slack state, or by simply subtracting their slack-state-sums from one another.
2. Sample the native basin less. Although these windows are not as expensive to run, they also show so little variability across both replicas and slices that having run them for much less time (a factor of ten less) would not have perturbed the free energy change estimates.
3. Extend the sampling for windows that show deviation across slices. By the same logic as the previous two points, windows that show high variability are necessarily more poorly defined, and will therefore change the resultant free energy change of stretching the most if more completely sampled. Use the standard deviation of the FEC derivative approach to identify which regions produce this variability and to determine the starting structures for follow up simulations that would target them.
4. Use window to window Hamiltonian replica exchange. Using replica exchange to exchange restraints between neighboring windows is a classic technique in umbrella sampling simulations that enhances convergence across the FEC.^44^ While our computational environment restricted us from using this technique on all windows, identifying poorly behaved windows given sequence length (for us the range is consistently 18-25 Å) may markedly speed up convergence in these regions.
5. Bias other reaction coordinates to more efficiently sample the loss of structure. End-to-end distance may not be the most efficient path between the native structure and a stretched state. Because such simulations possess both well-defined start and endpoints, path-finding methods like nudged elastic band^94–96^ or finite temperature string methods^97–100^ might provide more useful window spacings, window positions, or additional restraints. Such methods might yield reaction coordinates that, while specific to RNA (and likely to RNA of a particular length) are not particular to any sequence, allowing results from a few careful reaction coordinate finding simulations to be used for many new RNA sequences.
6. Use fixed positional restraints rather than distance restraints. This would eliminate the RNA’s translational and rotational rigid body degrees of freedom, restraining the end-to-end positions of the RNA would the allow a hexagonal prism or rectangular box to be used, which would reduce the number of solvent molecules in more extended windows. This also mimics stretching experiments more directly, since having both ends of the chain tethered prevents long axis rotations and, for a given extension distance, center of mass translations.
7. Analyze error by computing the average difference between RNA stretches and melts (Table 1). This value can be thought of as the sequence-independent free energy of relaxation from stretched to random coil states (and sequence independent errors of stretching or of the force field) that is being subtracted away in the thermodynamic cycle. This error term should contain both systemic, sequence-specific error and sampling error, but the variability in it is already low enough to discern outliers (GUAAUA is more overly stable than the other two sequences). Adding to the diversity of stretched sequences would make such an observation a more firm conclusion. It would also allow the approximate accuracy of a newly stretched sequence to be estimated without multiple replicas or reference to a pre-existing melt, which will be important in the future as this technique becomes more accurate and the collection of sequences with known reference melts to which it has been applied grows.

## Conclusions

In this study we furthered an approach to compare *in vitro* folding stabilities and *in silico* free energy curves. We refined methods for analyzing such data, and situated future studies on this topic to add sequence diversity more efficiently. Most notably, we showed that the thermodynamic landscape of FF99+bsc0+χ_OL3_ is chemically accurate where previously questions had lingered about whether differences could be due in part to lack of convergence.

## Acknowledgement

We thank Dr. Tod D. Romo for help writing analysis code using the software library LOOS. Dr. Jeffrey Zuber graciously provided us with the correlations between nearest neighbor parameters. This work was supported by grant NIH R01 GM076485 to D.H.M. and L.S. was a trainee of NIH T32 068411.

## Supporting Information Available

The following items are available as supplements to the text.

- hairpin-pulling-supplement.pdf: Tables of fit constants to curve segments.
- hairpin-pulling-analysis.zip: Scripts for analyzing free energy, free energy variance, and our raw distance files.

This material is available free of charge via the Internet at http://pubs.acs.org/.

## Graphical TOC Entry

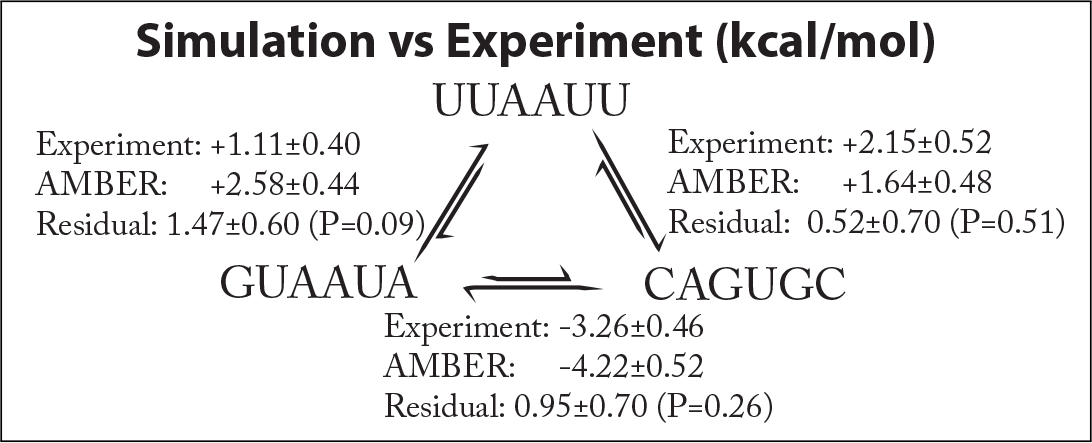

